# SCGclust: Single Cell Graph clustering using graph autoencoders integrating SNVs and CNAs

**DOI:** 10.1101/2025.01.28.635357

**Authors:** Teja Potu, Yunfei Hu, Rituparna Khan, Srinija Dharani, Jingchao Ni, Liting Zhang, Xin Maizie Zhou, Xian Mallory

## Abstract

Intra-tumor heterogeneity (ITH) is a compounding factor for cancer prognosis and treatment. Single-cell DNA sequencing (scDNA-seq) provides cellular resolution of the variations in a cell and has been widely used to study cancer progression and responses to drug and treatment. While the low coverage scDNA-seq technologies typically provides a large number of cells, accurate cell clustering is essential for effectively characterizing ITH. Existing cell clustering methods typically are based on either single nucleotide variations (SNV) or copy number alterations (CNA), without leveraging both signals together. Since both SNVs and CNAs are indicative of the cell subclonality, in this paper, we designed a robust cell clustering tool that integrates both signals using a graph autoencoder. Our model co-trains the graph autoencoder and a graph convolutional network (GCN) to guanrantee meaningful clustering results and to prevent all cells from collapsing into a single cluster. Given the low dimensional embedding generated by the autoencoder, we adopted a Gaussian Mixture Model to further cluster cells. We evaluated our method on eight simulated datasets and a real cancer sample. Our results demonstrate that our method consistently achieves higher V-measure scores compared to SBMClone, a SNV-based method, and a K-means method, which relies solely on CNA signals. These findings highlight the advantage of integrating both SNV and CNA signals within a graph autoencoder framework for accurate cell clustering. SCGclust is publicly available at https://github.com/compbio-mallory/cellClustering_GNN.

## Introduction

Cancer cells evolve by acquiring new mutations. Two important types of mutations are single nucleotide variation (SNV) and copy number alterations (CNA). A cancer sample often contains multiple subclones, each distinguished by a unique set of SNVs and CNAs. This diversity within a tumor is referred to intra-tumor heterogeneity (ITH).

ITH has confounded the cancer treatment, prognosis, and metastasis prevention (Lawrence et al., 2013; Burrell et al., 2013; Turajlic et al., 2019; Lawson et al., 2018). Specifically, ITH has been known to lead to heterogeneous cellular phenotypes that exhibit differential response to therapies, including the emergence of drug resistant cancer cells (Dagogo-Jack and Shaw, 2018; Marusyk et al., 2020).

Over the last decade, the advent of single-cell DNA sequencing (scDNA-seq) has revolutionized the study of ITH. This technology provides an unparalleled ability for characterizing ITH at the level of individual cells, enabling a deeper understanding of tumor complexity.

Given that each cell belongs to a subclone, with each subclone characterized by a unique set of single nucleotide variation (SNV) and copy number alteration (CNA), it is natural to cluster cells using SNV and/or CNA profiles, where each cluster corresponds to a subclone. However, this task is complicated by the technical limitations of single-cell DNA sequencing (scDNA-seq). The current cost-effective scDNA-seq technologies, such as degenerate oligonucleotide-primed PCR (DOP-PCR) (Carter et al., 1992; Navin et al., 2011; Baslan et al., 2012) and direct library preparation (DLP and DLP+) (Zahn et al., 2017; Laks et al., 2019), are designed to sequence hundreds or thousands of cells, each at a low cost. However, these methods typically produce very shallow coverage, often around 0.02X. This shallow coverage makes it challenging to cluster the cells since approximately 98% of the genomic region remains uncovered by a single read.

On the other hand, although multiple displacement amplification (MDA) (Hou et al., 2012; Wang et al., 2014) can produce high coverage scDNA-seq data, it is much more costly per cell than DOP-PCR or DLP/DLP+. Since the goal of cell clustering is to comprehensively characterize ITH, sequencing a larger number of cells increases the likelihood of capturing all subclones, thereby providing a more complete understanding of ITH. Therefore, in this paper, we focus on cell clustering for the scDNA-seq technologies capable of generating hundreds to thousands of cells, even at extremely low coverage.

In the past, despite that most of the SNVs lack informative signals due to insufficient coverage in such data, SNVs have still been used for cell clustering for the shallow scDNA-seq. Specifically, SBMClone is an SNV-based method that employs a stochastic block model (SBM) for accurate cell clustering. Nevertheless, SBMClone does not incorporate CNA signals into cell clustering. To date, no method exists that considers both SNV and CNA signals together for cell clustering in the shallow scDNA-seq data. We reason that since both SNVs and CNAs arise during cancer cell evolution, combining these two signals together for cell clustering can produce better results.

Recently, graph autoencoders have been widely used in genomic sequencing analysis Hu et al. (2024). For example, ADEPT (Hu et al., 2023) leveraged a graph autoencoder to learn the low-dimensional latent embedding of each spot for spatial transcriptomics data, which was then used to cluster spatial spots. ADEPT modeled each spot as a node in the graph, with its gene expression data serving as the node feature. The spatial context of the spots was represented by the presence or absence of edges in the graph. By utilizing both the node features and edge weights, ADEPT effectively integrated gene expression signals with spatial context. Inspired by ADEPT’s design, we developed SCGclust and compared it against SBMClone and a K-means-based method that uses only CNA signals across eight simulated datasets and a real breast cancer dataset. We found that SCGclust consistently outperformed SBMClone and the CNA-based method, producing more accurate and robust clustering results. Additionally, SCGclust demonstrated greater resilience to extreme cases and was less sensitive to the lack of or wrong signals.

## Method

The proposed SCGclust framework integrates graph-based learning with clustering methods to analyze genetic variation data at a single-cell resolution. It employs a graph neural network (GNN) architecture to model relationships between cells based on genomic features and utilizes advanced clustering techniques to partition cells into distinct subgroups.

**Fig. 1** shows an overview of the SCGclust framework. SCGclust takes as the input a cell-by-SNV matrix and a read count matrix containing the CNA signals (**Fig. 1A**). Using these inputs, it builds a graph where each node represents a cell, the node feature corresponds to the CNA signals, and the edge weights between two nodes reflect the similarity between SNV profiles of two cells (**Fig. 1B**). Given the graph built in **Fig. 1B**, SCGclust constructs a graph autoencoder consisting of two encoder layers (layers 1 and 2) followed by two decoder layers (layers 3 and 4). This architecture in **Fig. 1C** reduces the dimensionality of the node features while accounting for node features of neighboring nodes in the graph, weighted by the edge weight. SCGclust co-trains the parameters in the graph autoencoder and a two-layered graph convolutional network (GCN) shown in **Fig. 1D**, the latter of which takes the low dimensional embedding from the graph autoencoder as the node feature. The objective function is composed of three terms, the reconstruction mean squared error (MSE) term from the graph autoencoder, the modularity and the collapse regularization term from GCN. Finally, SCGclust clusters the cells based on the latent features extracted from the low-dimensional representation of the last encoder layer, applying an existing clustering method Gaussian Mixture Model (GMM) (**Fig. 1E**). In the following, we describe in more detail the graph construction, graph autoencoder architecture, co-training graph autoencoder and GCN, and the clustering step.

**Figure 1:**
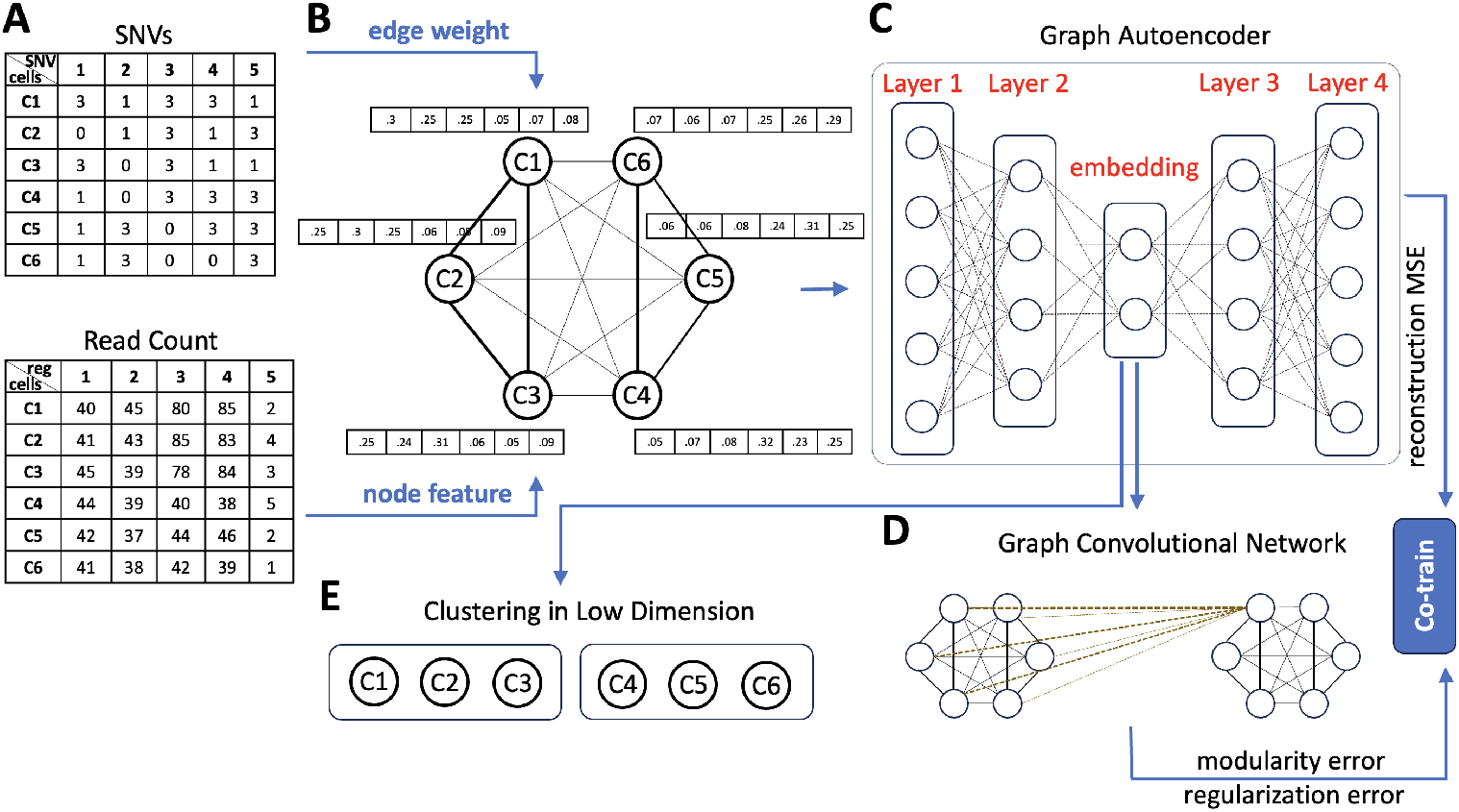
Overview of SCGclust. **A**. There are two inputs to SCGclust, the cell by SNV matrix (top) and the cell by genomic region matrix (bottom). The cell by SNV matrix has entries “1”, “0” and “3”. The “1” and “0” entries represent that the SNV is present and absent in the cell, respectively. “3” entries represent that there is no read covering the site and thus the signal is missing. The cell by genomic region matrix has the read count for each genomic region at each cell. Six cells (C1-C6) are shown as an illustration. It can be observed that C1-C3, and C4-C6 have relatively similar SNV and CNA profiles, respectively. **B**. The two matrices will then be used as the edge weight and the node feature for the graph autoencoder. The graph autoencoder has six nodes, representing the six cells. On top of each node is a vector of node feature, which uses the cosine similarity vector of the read count that reflects the CNA signal. Between every two nodes is an edge weight, represented by the dot product of the SNV profiles between the two cells. Here C1, C2 and C3 have larger edge weights (thicker edges) because their SNV profiles are closer to each other. Similarly, C4, C5 and C6 have larger edge weights (thicker edges). **C**. The built graph will be the input for the graph autoencoder, which will reduce the dimension of the node features in the encoder, and recover the original node features in the decoder. The dimension reduction process also considers the edge weight such that the two cells having similar SNV profiles will have more similar embedding in the low dimension. The graph autoencoder in total has four layers, layers 1 and 2 are the encoder, and layers 3 and 4 are the decoder. **D**. A Graph Convolutional Network (GCN) is co-trained with the graph autoencoder, with the objective function composed of three terms, the reconstruction mean squared error (MSE) term, the modularity term and the collapse regularization term. **E**. Finally, we perform the cell clustering based on each cell’s embedded low dimension from the graph autoencoder using Gaussian Mixture Model.

### Graph Construction

To represent the genetic variation data as a graph, we construct an adjacency matrix and define node features, enabling GNN-based learning.

#### Edge weight adjacency matrix

The edge weight adjacency matrix is computed based on the dot product of the SNV profiles, which is used as the edge weights in the graph. This matrix quantifies pairwise similarities between all cells, with each entry ranging from 0 to 1, where 0 indicates no similarity and 1 indicates identical profiles. Due to that a majority of the entries in the SNV profile consist of missing entries (“3”), we replace all missing entries with “0”s before calculating the dot product of every two SNV profiles, following the same practice of SBMClone (Myers et al., 2020). In this way, the missing entries are treated the same way as the absence of the SNV. In fact, even if a genomic site is covered by a reference-supporting read but not by a variant-supporting read, this site might still harbor an SNV if it lacks sufficient read coverage. Conversely, if a site is covered by a variant-supporting read, it is highly likely that an SNV is present, given that the false positive (FP) rate in scDNA-seq is relatively low (0.01-0.05). Therefore, if a site is covered only by a reference-supporting read, we cannot definitively determine whether the site harbor an SNV, as the variant-supporting read may not have been sampled or sequenced. In our simulated dataset section, such an uncertainty is also reflected in the simulated high false negative (FN) rate. Due to the same reason, we use dot product instead of other alternative metrics. When two cells have “0” and “1” on the same SNV site, we cannot be certain whether they are truly different, since the “0” could be the result of insufficient sequencing of variant-supporting reads rather than the true absence of a variant. However, when both cells have “1”s at the same site, we can confidently infer that they are similar at that site. The dot product captures this confidence by emphasizing shared SNV calls between cells.

#### Node Features

Node features are derived from the raw read counts of each cell, reflecting the CNA signals. However, instead of directly using the raw read count as the node features, we calculated the normalized cosine similarity between the raw read count vector of a cell and those of all other cells, resulting in a vector of normalized cosine similarities for each cell. This vector is used as the node feature. This approach avoids directly relying on raw read counts, which are prone to fluctuations caused by sequencing artifacts. Additionally, it eliminates dependence on the absolute values of raw read counts, which vary with the sliding window size and genome coverage. In contrast, cosine similarity is invariant to absolute values and more robust to fluctuations in read counts unrelated to CNAs. While it is possible to use inferred CNAs instead of raw read counts, we opted not to do so in this study to avoid errors introduced during CNV inference. Nevertheless, using inferred CNAs as the node feature has its advantage, and these trade-offs are discussed further in the conclusion.

### Graph Autoencoder Architecture

The SCGclust framework employs a Graph Attention Autoencoder to learn the latent representations of node features. The architecture consists of two main components: an encoder and a decoder, leveraging Graph Attention mechanism with Graph Convolutional Networks (GCNs).

#### Encoder and Decoder

Both the encoder and decoder comprise two Graph Attention Convolution layers. The encoder transforms input features (called **X**) and adjacency matrix (called **A**) into latent representations (called **h**^(2)^), while the decoder reconstructs the original input from these latent features. **Eq. 1** shows the mathematical equation of this process.

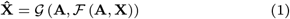

where ℱ and 𝒢 denote the encoder and decoder functions, respectively.

To enhance consistency between the encoder and decoder, weight sharing is employed. Let **W**^(*l*)^ denote the weight of the *l*-th layer, the weights of corresponding layers are tied via transposition, as shown in **Eq. 2**.

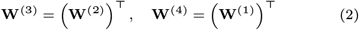

We train the encoder and decoder layers and learn the weights such that an objective function is minimized. One of the terms in our objective function is the reconstruction MSE between the input feature (**X**) and the recovered feature from the decoder 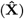, shown in **Eq. 3**. Such a term, combined with the modularity term and the regularization term shown in **Eq. 11**, will constrain the weights in the graph autoencoder.

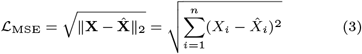

#### Attention Mechanism

In this section, we describe how we obtain the embedding **h**^(2)^ while engaging the edge weight adjacency matrix. Our graph autoencoder operates by allowing each node to attend to its neighbors through learned attention coefficients. In more detail, each node *i*’s feature vector **x**_*i*_ is first projected into a new representation **z**_*i*_ via **W**^(1)^:

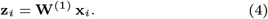

For every pair of nodes *i* and *j*, we compute a raw attention score 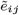 by applying a learnable vector **a** to the concatenated transformed feature [**z**_*i*_ ∥ **z**_*j*_]:

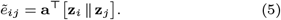

To respect the graph structure which contains the edge weight, we multiply each raw score by the corresponding adjacency entry *A*_*ij*_ to create weighted attention score:

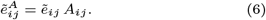

We then apply a LeakyReLU activation to obtain the final attention logits:

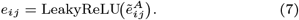

Next, these logits are normalized across nodes via a softmax, ensuring that the attention coefficients sum to 1 for node *i*:

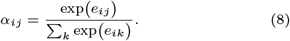

Finally, each node aggregates its neighbors’ features weighted by these learned attention coefficients:

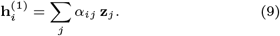

Here 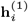 represents the hidden feature for node *i* from layer 1 that incorporated node *i*’s original feature **x**_*i*_ and the edge weight adjacency matrix **A**. This whole process generates a context-aware feature update by calculating the attention coefficient *α*_*ij*_ for each node *j*, and it allows graph autoencoder layers to learn which neighbors are most important for each node’s representation.

Notice that **h**^(1)^ is the hidden feature learned from layer 1 for node *i*. The second layer’s output, **h**^(2)^, is simply **W**^(2)^ **h**^(1)^ without an application of the attention weights. **h**^(2)^ is the latent embedding that will be fed to the Graph Convolutional Network (GCN) in the next step.

### Co-training Graph Autoencoder and Graph Convolutional Network (GCN)

(Tsitsulin et al., 2023) introduced a GCN that has been designed specifically for clustering in which the objective function contains both a modularity term which encourages meaningful clustering and a collapse regularization term which discourages from all cells collapsing into one cluster. We adopt its model and combine it with our graph autoencoder, such that the objective function includes not only the reconstruction MSE error, but also a spectral modularity term and a regularization term. In more detail, we first normalize the adjacency matrix **A** to be 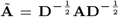, and we use **Ã** as the edge weight in a two-layer GCN that is operated on the embedding **h**^(2)^ from the graph autoencoder (**Fig. 1D**). We then apply a softmax to the output of GCN to obtain the soft cluster assignment **C**:

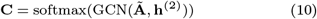

Given the soft assignment **C** from GCN, we then aim to minimize the following objective function,

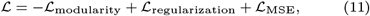

in which 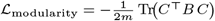, and 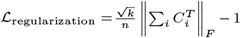. Here *C* ∈ ℝ^*n×k*^ is the soft cluster assignment matrix, 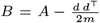 (with *A* the adjacency matrix and *d* the degree vector), *k* is the number of clusters, *m* =Σ_*i*_ *d*_*i*_*/*2 is the number of edges, and *n* is the number of nodes. ∥·∥_*F*_ represents the Frobenius norm. ℒ_MSE_ has been shown in **Eq. 3**. The first term (modularity) encourages meaningful clusters in the graph; the second term (collapse regularization) discourages all nodes from collapsing into a single trivial cluster; and the third term ensures the reconstructed feature is as close to the input feature as possible in the graph autoencoder.

The weights and parameters in both graph autoencoder and GCN are trained together to minimize *L* in **Eq. 11**. This combined objective function enforces high-quality clusters while preserving relevant node feature reconstruction.

### Gaussian Mixture Model for Clustering

Final cluster assignments are obtained by applying Gaussian Mixture Model (GMM) clustering on the learned embeddings **h**^(2)^ from the graph autoencoder. GMM models the latent space as a mixture of *k* Gaussian distributions, assigning each node to a cluster based on its learned representation. This two-step process leverages GCNs for high-level feature extraction and GMM for refined clustering, ensuring robust and interpretable partitioning of graph-structured data. Here *k* is given by the user. We discuss in the future work the implications of selecting *k* and a future work to automate the selection of *k*.

## Results

### Simulated Dataset

We systematically benchmarked SCGclust’s performance by varying eight key variables: the number of subclones, number of cells, number of SNVs, number of CNAs, false positive (FP) rate of SNVs, false negative (FN) rate of the SNVs, missing rate of the SNVs, and the noise level on the CNA signal. The eight variables, along with their values and default values, are listed in **Table 1**. For each experiment, we varied one variable while keeping all others at their default values. To ensure robust results and minimize the impact of random fluctuations, each setting was repeated five times. Among these variables, the missing rate of the SNVs is the most critical to our study, ranging from 0.95 to 0.98 and 0.99, with 0.98 set as the default. These missing rates correspond to coverage of 0.05X, 0.02X, and 0.01X, respectively.

**Table 1.** Summary of simulated datasets. Each row represents a simulated dataset with an index (first column) whose values of the varying variable (second column) are shown in the third column. The default value is denoted by “(d)” on its right.

|  |  |  |
| --- | --- | --- |
| 1 | Number of subclones | 2, 4 (d), 6 |
| 2 | Number of cells | 100, 500 (d), 1000 |
| 3 | Number of SNVs | 0, 1000, 5000 (d), 10000 |
| 4 | Number of CNAs | 100, 200 (d), 300 |
| 5 | False Positive Rate | 0.01, 0.05 (d), 0.1 |
| 6 | False Negative Rate | 0.55, 0.6 (d), 0.65 |
| 7 | Missing rate | 0.95, 0.98 (d), 0.99 |
| 8 | CNA noise | 0.3, 0.5(d), 1 |

In the following, we describe our simulator in detail, followed by a benchmark of SCGclust on the simulated dataset.

### Simulator

To mimic the whole process of cancer evolution, we employed the Beta-splitting model to generate a phylogenetic tree, following the practice of SCsnvcna (Zhang et al., 2023). The process begins with the generation of a clonal tree, where each leaf node represents a subclone of cells. The number of subclones is predetermined, and the tree structure is built using a Beta-splitting model which stochastically determines the percentage of each subclone.

In more detail, the simulation starts with a normal root node. Each node that has not split yet has a chance to split into two sub-clones. This process goes on recursively until the desired number of leaf nodes is reached.

CNAs are then placed on the edges based on the branch lengths, with random intervals selected across chromosomes. The branch length of each edge is determined according to an exponential distribution, following the practice of (Mallory et al., 2020). Each CNV corresponds to a randomly selected genomic region that is either amplified or deleted. After determining the copy number states, we simulate the read count following the practice of (Zhang et al., 2024). In more detail, we use normal cells from the T10 dataset (Navin et al., 2011) to generate corresponding read counts. Noise is introduced into the copy number reads to simulate real-world biological variability. Specifically, 50% of the data rows for each set of leaf cells are randomly selected using a binary mask. Noise is then added to the selected copy numbers using a normal distribution with a mean of zero and a specified standard deviation, reflecting measurement fluctuations. Negative copy numbers are constrained to zero to maintain biological plausibility. The read counts for each cell are adjusted based on the noisy copy number values to ensure consistency with the original dataset while capturing the introduced variability.

SNVs are also placed on the edges according to branch length, with their positions randomly sampled to ensure no overlap with CNAs. Cells are assigned to the leaves based on their cell proportions. We then walk the path from the root to the leaf, and the SNVs and CNAs that are placed on the edges of the path are assigned to the cells corresponding to the leaf. In this way, we generate a complete genotype matrix. To simulate real-world conditions, we introduce false positives, false negatives, and missing data to the genotype matrix, given the variable used in each simulation setting. It is worth to note that we intentionally set the FN rate to be above 0.5, because when a site is covered by reads, most of the time it is covered by only one read. For a heterozygous SNV that falls in a copy number neutral area, half of the chance that the read is a reference-supporting read. Thus the false negative rate shall include both the low coverage factor and the sequencing artifact in the scDNA-seq. We set the FN rate to range between 0.55 and 0.65 in our simulator. For the missing rate, we set it to be 0.95, 0.98, and 0.99, reflecting the low coverage of scDNA-seq at 0.05X, 0.02X, and 0.01X, respectively.

### Benchmarking results

We evaluated SCGclust, SBMClone (Myers et al., 2020), and a K-means-based method that uses only the read count on simulated datasets. Since this K-means-based method uses the CNA signals, we refer to it as “Kmeans-CNA”. SCGclust was tested in two modes: one mode where silhouette score was used to select the best epoch without knowing the ground truth, and another as a reference where the ground truth was assumed to be known for selecting the best epoch. While the ground truth is not available in real-world applications, this reference mode helps determine whether any epoch achieves a good clustering result, providing a benchmark for assessing the effectiveness of SCGclust and SBMClone.

We compared four methods on the simulated datasets, SCGclust using silhouette score to select the optimal epoch (referred to as “SCGclust-silhouette”), SCGclust using the ground truth to select the optimal epoch (referred to as “SCGclust-GT”), SBMClone, which uses only SNV signals, and Kmeans-CNA, which relies solely on CNA signals. In particular, we compared SCGclust with SBMClone and Kmeans-CNA to investigate whether SCGclust, which integrates both SNV and CNA signals, outperforms methods that rely soley on either SNVs or CNA signals but not both. Given its ability to combine both signals, SCGclust is expected to have a distinct advantage.

We plotted the V-measures of these four methods in **Fig. 2**. V-measure is a metric that measures quantitatively the clustering result. It assesses the balance between homogeneity and completeness, in which homogeneity measures how much each inferred cluster contains only the cells in a single true subclone, and completeness measures how well all members of a true subclone are assigned to the same inferred cluster. V-measure ranges from 0 to 1, whereas a score of 1 indicates the perfect clustering, and 0 indicates poor clustering. It can be observed that SCGclust’s V-measures are predominantly around and above 0.6 across all eight variables, whereas SBMClone and Kmeans-CNA generally have V-measures around or below 0.6. The only instance where SBMClone achieves a V-measure well above 0.6 is when the missing rate is as low as 0.95 (**Fig. 2G**), with V-measures ranging between 0.7 and 0.8. Looking into details of the performance of these methods, we found that all methods perform better as the number of SNVs increases (**Fig. 2C**). This result is expected, as more SNVs provide greater signal strength and clustering power.

**Figure 2:**
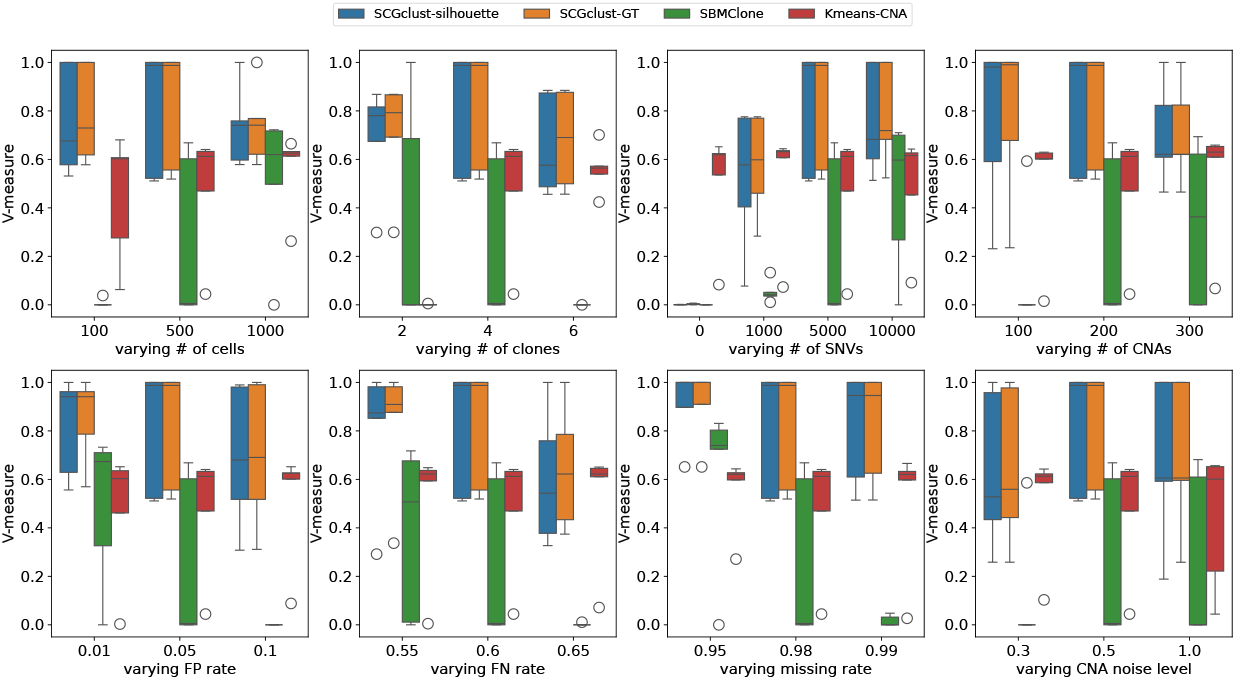
Boxplots are shown for V-measure for SCGclust using silhouette score to select the best epoch (SCGclust-silhouette, in blue), SCGclust using the ground truth clustering to select the best epoch as a reference (SCGclust-GT, in orange), SBMClone (SBMClone, in green) and a K-means method that uses the read count which is a CNA signal to cluster the cells (Kmeans-CNA, in red). Eight variables were varied to study the performance of these four methods, including **A**. number of cells, **B**. number of clones, **C**. number of SNVs, **D**. number of CNAs on the top panel, and **E**. FP rate, **F**. FN rate, **G**. missing rate, and **H**. CNA noise on the bottom panel. Median of each boxplot was highlighted with the black horizontal line. Each set of variables was repeated for five times to avoid extreme cases.

To assess whether SCGclust can function effectively when only CNA signals are provided as the node features, and the edge weight does not reflect the SNV signal, we conducted an additional test. Specifically, in **Fig. 2C**, we set the SNV number to zero, assigned a very small constant weight (0.001) to all edges, and tested SCGclust’s performance under this condition. We found that under this setting, SCGclust fails to render meaningful clustering results, indicating that SCGclust’s result does benefit from SNV signals. Thus although weak, these SNV signals provide valuable information for clustering. It is also noticeable in **Fig. 2C** that when the SNV count is as low as 1,000, SBMClone’s V-measure drops to near zero, showing that SBMClone requires a large number of SNVs for effective clustering. In contrast, SCGclust shows no significant decline in V-measure as the number of SNVs decreases, maintaining a median V-measure of around 0.6 with only 1,000 SNVs.

Furthermore, in addition to the SNV number, we observed that SBMClone is much more sensitive to variations in the number of cells (**Fig. 2A**) and the number of clones (**Fig. 2B**) than SCGclust. Specifically, when the number of cells is as low as 100 or the number of clones is as high as 6, SBMClone’s V-measure drops to around zero. In contrast, SCGclust maintains a V-measure above 0.5 in both scenarios. Additionally, we noticed that all methods, except Kmeans-CNA, perform worse as the SNV missing rate increases (**Fig. 2G**). This outcome is expected because a higher SNV missing rate renders less signal for clustering. Notably, SBMClone’s V-measure drops near zero when the SNV missing rate is at 0.99, showing that SBMClone is very sensitive to the SNV missing rate.

Since SBMClone is a method based on SNV only, it is expected that the absence of a strong SNV signal would lead to poor performance. In contrast, although SCGclust’s V-measure decreases with the increase of the SNV missing rate, it remains above 0.5 even at the highest SNV missing rate of 0.99. This highlights the advantage of SCGclust, which integrates both CNAs and SNVs. A similar pattern is observed with varying SNV FP and FN rates (**Fig. 2E, F**). While Kmeans-CNA’s performance is stable, SCGclust and SBMClone exhibit declining V-measure as the SNV FP and FN rates increase. However, the decrease in SCGclust’s V-measure is less pronounced compared to SBMClone, the latter of which drops to near zero across all five repetitions when the SNV FP rate is as high as 0.1 or the SNV FN rate reaches 0.65. This again shows that SBMClone is more sensitive to SNV FP and FN rate than SCGclust. Furthermore, we expected an increase in the V-measure for SCGclust and Kmeans-CNA with an increased number of CNAs, as both methods leverage the CNA signal. However, such a trend is not obvious in the plot (**Fig. 2D**), suggesting that even a low CNA number of 100 is sufficient for cell clustering.

Lastly, we investigated the effectiveness of the silhouette score in selecting the optimal epoch by comparing SCGclust-silhouette and SCGclust-GT. Our results show that SCGclust-silhouette (in blue bars) closely aligns with SCGclust-GT (in orange bars) in most cases, demonstrating that the silhouette score is a reliable indicator of clustering performance and it effectively guides the selection of the best epoch.

In summary, SCGclust outperforms both SBMClone and Kmeans-CNAs, and SCGclust is much less sensitive to the signal of SNVs than SBMClone, showing the advantage of integrating both SNV and CNA signal. SCGclust’s V-measure can reach values exceeding 0.9 even when at the missing rate of 0.95, which corresponds to a coverage of 0.05X.

### Real Dataset

We further tested SCGclust on a real breast cancer sample T10, which has 100 cells and the Fluorescence Activated Cell Sorting (FACS) indicating the presence of four distinct clusters, D, H, A1 and A2 (Navin et al., 2010). The FACS result can serve as a ground truth to evaluate SCGclust’s performance. The average coverage of T10’s data is about 1.81X. We used SCcaller (V2.0.0), a single-cell mutation caller for scDNA-seq data, to detect SNVs on each cell. In total, there are 48,898 SNVs detected for all cells. We then applied a filter for variant allele fraction *>* 0.03, selected only nonsynonymous SNVs in exonic regions, and obtained a total of 3,105 SNVs. These 3,105 SNVs were the input to SCGclust and SBMClone. For CNA signals, we used read counts for a sliding window of 500kbp for each cell. This CNA signal was the input to both SCGclust and Kmeans-CNA. We compared the performance of SCGclust-silhouette, SCGclust-GT, SBMClone, and Kmeans-CNA, with their V-measures (using the FACS results as the ground truth) being 0.61, 0.64, 0.037, and 0.22, respectively. SCGclust has a much better performance than SBMClone and Kmeans-CNA. We then investigated whether SBMClone’s performance could improve by using all 48,898 SNVs. Our results show that while SBMClone’s performance did improve slightly with more SNVs, the increase was limited, with the V-measure rising to 0.077. We reason that since in the simulated data, SBMClone’s V-measure was near zero when the cell number is as low as 100, it is expected that SBMClone exhibits similarly low V-measure in the T10 dataset with the same cell count. Thus, the large number of SNVs cannot compensate for the low number of cells for SBMClone. In contrast, SCGclust is robust to variations in cell number, maintaining a V-measure above 0.6 even with only 100 cells.

It is worth to note that in this real dataset, the silhouette score can be used as a metric to select the best epoch since the V-measures between SCGclust-silhouette and SCGclust-GT are close to each other. This indicates that even when SCGclust is applied to a real dataset without a ground truth clustering, the silhouette score should consistently be able to select the optimal epoch. Since in practice, we are not given the true cluster number, we then investigated the performance of SCGclust when the given cluster number is slightly off. When setting the cluster number to be 3, SCGclust’s V-measure is 0.6619 for SCGclust-silhouette, and 0.6688 for SCGclust-GT, showing an even higher V-measure than the case when the correct cluster number is given. This is likely due to that the A2 subclone has only four cells, and separating these four cells to be a stand-alone subclone is challenging. When setting the cluster number to be 5, SCGclust’s V-measure is 0.33 for SCGclust-silhouette, and 0.49 for SCGclust-GT, showing a drop of V-measure when the cluster number is one off. However, the resulting V-measure is still significantly higher than SBMClone and Kmeans-CNA.

## Conclusion

ITH has been known to confound cancer treatment. To characterize ITH, cell clustering is the first and foremost step in scDNA-seq data. In this paper, we presented SCGclust, a cell clustering method that is applicable to shallow scDNA-seq data integrating both the SNV and CNA signals. SCGclust adopts a graph autoen-coder framework, in which each node represents a cell, the node feature is the read count reflecting the CNA signal, and the edge weight is the SNV similarity between two cells. SCGclust co-trains the graph autoencoder and a GNN with an objective function that guarantees optimal clustering result. By integrating both SNVs and CNAs, SCGclust outperforms the methods that rely on either signal but not both, including SBMClone and Kmeans-CNA, a K-means algorithm based on read count only. We tested SCGclust, SBMClone and Kmeans-CNA on eight simulated datasets and one real dataset. We found that SCGclust has an advantage of being robust in the extreme cases such as very low coverage reflected by a high missing rate, low number of cells, and high SNV FP and FN rates. In particular, SCGclust’s V-measure is mostly consistently above 0.6, whereas SBMClone’s V-measure drops to zero in several extreme cases. SCGclust has also been shown to have a higher V-measure than SBMClone and Kmeans-CNA on the real dataset T10. These show that SCGclust is a robust tool that can be used to cluster the low coverage scDNA-seq data.

Regarding potential improvement on SCGclust, since SCGclust does not rely on the fact that the copy numbers are integers but uses directly the read count instead of the inferred copy number, a future direction is to incorporate either the inferred absolute copy numbers instead of the read count, or combining the CNA inference and cell clustering together in a more complex model. Nevertheless, the current model that uses the read count directly is still advantageous in avoiding the CNA inference error that may be passed by a CNA caller, which is still challenging in scDNA-seq data (Zhang et al., 2024).

Identifying the correct cluster number is non-trivial to all cluster algorithms. In our current setting, we assume that the cluster number is known. We further test SCGclust’s performance when given a wrong cluster number on the real dataset. SCGclust’s V-measure drops from 0.61 to 0.33 when the given cluster number is one higher than the correct number, but increase from 0.61 to 0.66 when the given cluster number is one smaller than the correct number. Even though provided with the wrong cluster number, SCGclust’s V-measures are still higher than that of SBMClone’s (0.037) and Kmeans-CNA’s (0.22) when they are provided with the correct cluster number. A future direction is to automate the cluster number selection so that the users do not have to input the cluster number to SCGclust.

## Author contributions statement

X.M., X.M.Z., and L.Z. conceived and supervised the project. T.P., Y.H., J.N., L.Z., X.M.Z, and X.M. designed the model, T.P., Y.H., S.D., L.Z., and R.K. conducted the experiments, and T.P., L.Z., Y.H., X.M.Z, and X.M. wrote the manuscript. All authors reviewed and approved the manuscript.

## Acknowledgments

The authors thank the anonymous reviewers for their valuable suggestions. This work is supported in part by funds from the NIH NIGMS Maximizing Investigators’ Research Award (MIRA) R35 GM146960.

